# Designing Well-Organized Touch-Spun 3D Scaffolds: The Role of Interfiber Spacing in Cell Proliferation

**DOI:** 10.64898/2026.09.16.752205

**Authors:** Kristina Peranidze, Nataraja S. Yadavalli, Mohammad Aghajohari, Sergei Makaev, Mikhail Parker, Sergiy Minko, Vladimir Reukov

## Abstract

Three groups of touch-spun 3D scaffolds with uniform fiber diameters and varying interfiber spacing were studied to identify optimal conditions for NIH/3T3-GFP fibroblast growth. All scaffold groups exhibited higher cell viability relative to control levels after 14 days of culture, with the highest viability observed for scaffolds with a 50.61 ± 15.35 μm interfiber spacing. Although cell counts indicated comparable cell growth across groups, scaffolds with medium and large interfiber spacing demonstrated an improved capacity to sustain viable cells without necrotic core formation. These findings provide insight into the optimal structural design of touch-spun 3D scaffolds for tissue engineering applications.

## 1. Introduction

The fabrication of three-dimensional (3D) scaffolds with optimized architectures for efficient cell proliferation has been a central focus of joint research in materials science and cell biology for over two decades. As the native extracellular matrix (ECM) of tissues primarily consists of complex fibrous protein networks [1], the development of polymer-based fibrous scaffolds with ECM-like features has become a major trend in biomimetic approaches to tissue regeneration [2]. The ECM is a tissue-specific parameter; therefore, the structural characteristics of fabricated fibrous scaffolds, such as fiber diameter, alignment, mesh size, crystallinity, and mechanical strength, must be tailored to the requirements of the target tissue.

The majority of tissues do not display a specific arrangement of ECM fibers into defined nano- and microfiber patterns. Accordingly, the skin dermis [3], loose connective tissue [4], adipose tissue [5], as well as parenchyma of organs [6, 7], generally lack a predominant global fiber orientation and are composed of irregular, mesh-like networks of collagen and elastin fibers. Therefore, when designing polymeric 3D scaffolds for these tissues, it is unnecessary to tune fiber arrangement within the cell-culturing platform. However, some tissues critically depend on the specific organization of protein fibers, as it underlies their physiological function. Examples include tendons and ligaments, where highly parallel collagen fibers enable efficient uniaxial load bearing [8]; skeletal [9] and cardiac muscle tissues [10], where aligned ECM fibers facilitate coordinated contraction; the cornea, with collagen fibers organized in highly ordered lamellae that confer both optical transparency and mechanical strength [11]; and blood vessels, where collagen and elastin fibers exhibit circumferential and longitudinal orientations to withstand pulsatile pressure [12]. Special attention has been given to peripheral and central nervous tissues, in which ECM fiber alignment directs axonal growth and supports regeneration. Previous studies [13–15] demonstrated that aligned fibers enhance neurite outgrowth of primary cortical neurons and PC-12 cells, promoting effective spinal cord and peripheral nerve repair. In line with these findings, our research group has shown that well-organized nanofiber scaffolds influence mouse neuroepithelial (NE-4C) cell behavior, guiding neurite extension along the fibers [16, 17]. Collectively, these findings underscore the importance of well-aligned fiber patterns for the regeneration of specific tissue types.

In addition to fiber alignment, a critical parameter in overall fiber arrangement is the spacing between fibers, commonly referred to as pore or mesh size. Interfiber spacing plays a key role in regulating nutrient supply from the culture medium and enabling efficient cell attachment during seeding [18]. For ECM-producing cell lines [19, 20], it also determines the area available for ECM deposition. Although the optimal spacing varies by cell type, several studies indicate that for mammalian cells with average diameters of ~10-20 μm, effective interfiber spacing typically ranges from ~40 to 100 μm [21–23].

Mechanical fiber-drawing techniques [24–26] have emerged as a particularly promising approach among the methods commonly employed to fabricate aligned fiber patterns with precise control over interfiber spacing. Unlike conventional electrospinning and its modifications, this relatively recent approach utilizes precise motorized systems for controlled fiber drawing and does not rely on the dielectric properties of the polymer solution, thereby expanding the range of materials suitable for scaffold fabrication. A notable example is the touch-spinning technique, which enables the production of well-aligned nano- and microfiber arrays with consistent interfiber spacing as small as a few microns [25]. This precision is achieved through programmable motor systems that coordinate both rotational motion of spinnerets and translational (Z-axis) motion of collecting frames. Furthermore, touch-spinning can be integrated with additive manufacturing technologies, facilitating the assembly of fiber arrays into complex, multilayer 3D scaffolds. The efficiency of touch-spun scaffolds has been reported for culturing several cell types, including mouse neuroepithelial cells (NE-4C) [16, 17], NIH/3T3 mouse fibroblasts [27], and mouse myoblast cells (C2C12) [28]. A recent study [29] further explores strategies to navigate the proliferation profile of insulin-producing INS-1 cell cultures within well-aligned touch-spun biomimetic scaffolds for tissue engineering applications.

In this study, we investigate the regularities of cell growth within well-aligned 3D microfiber scaffolds fabricated by integrating touch-spinning with additive manufacturing, with a particular focus on the role of interfiber spacing. The interfiber distance was precisely controlled by adjusting the translational speed of the fiber-collecting frame. GFP-labeled NIH/3T3 fibroblast cells were selected as a model cell system due to their robustness and ability to deposit ECM between adjacent fibers during proliferation. The performance of touch-spun scaffolds with varying interfiber spacing was evaluated through cell growth visualization, viability assays, and cell counting conducted at the end of the culture period. Through this approach, the work aims to identify the optimal interfiber spacing in touch-spun 3D scaffolds that promotes efficient cell attachment and growth.

## Materials and Methods

### Touch-spinning and Scaffold Assembly

Polycaprolactone (PCL; M_n_ = 80 kDa; Sigma-Aldrich), polyethylene oxide (PEO; M_w_ = 5,000 kDa; Sigma-Aldrich), and chloroform (purity ≥ 99.8 %; Thermo Fisher Scientific) were used to prepare blend solutions for touch-spinning. The blends contained PCL and PEO at weight fractions of 5 wt% and 0.25 wt%, respectively. Microfiber layers with uniform fiber diameters and well-controlled interfiber spacing were fabricated using a modified touch-spinning technique (CytoNest, Inc.) as described in the study [24]. Briefly, this technique employs the rotational motion of a spinneret that comes into contact with a solution droplet and stretches it around a collecting frame, while the Z-axis motion of the frame defines the spacing between adjacent fibers. Within the framework of this research, the Z-axis speed of the frame was adjusted to achieve interfiber spacings of 20 (Group 1), 40 (Group 2), and 60 μm (Group 3).

The produced fiber layers were assembled into four-layered scaffolds using customized 3D-printed grid spacers and outer frames, as described in our recent work [30]. Each layer of the final scaffold consisted of two orthogonally aligned touch-spun fiber sheets. The dimensions of the spacers were tailored to fit 6-well tissue culture plates (TCPs) with a well diameter of 35 mm, ensuring convenience of cell culture experiments.

### Characterization of Fiber Patterns

Fiber arrangement and alignment were assessed in orthogonally aligned touch-spun layers using an optical microscope (Olympus BX51; Evident Scientific, Inc.) equipped with a digital camera (Nikon D5300; Nikon Corp.). Images were captured at 5× and 20× to assess fiber organization. Fiber diameter and interfiber spacing were measured using Fiji software.

Fiber surface wettability was evaluated using the water contact angle method. Droplets of deionized water were placed on the surface of the touch-spun fiber sheets, and images of the droplets were captured using a CMLN-13S2M-CS digital camera (Point Grey Research, Inc.) connected to an optical microscope. Contact angles were quantified using ellipsoidal fitting in Fiji software with the Contact Angle Plugin.

### Cell Culture and Visualization

Prior to seeding onto scaffolds or control wells, NIH/3T3-GFP fibroblasts (Cell Biolabs, Inc.) were grown to moderate confluency in a T75 tissue culture flask containing DMEM supplemented with 20 % FBS and 2 % antibiotic-antimycotic solution. Culture was maintained in a humidified incubator at 37 °C with 5 % CO_2_, and the medium was replaced every other day. For cell preparation, the culture medium was aspirated, and the cells were rinsed twice with PBS. Cells were then detached by adding 0.25 % trypsin-EDTA solution and incubating the flask at 37 °C for 3-5 minutes under 5 % CO_2_. The resulting cell suspension was neutralized with fresh culture medium and transferred to a 15 mL centrifuge tube, which was spun at 1,500 rpm for 5 minutes to obtain a cell pellet. The supernatant containing trypsin was carefully removed using a glass pipette connected to a vacuum pump, and the pellet was resuspended in an appropriate volume of fresh medium. A 100 µL aliquot of the suspension was taken for cell counting.

Before cell seeding, multilayer scaffolds were subjected to high-power air plasma treatment (30 W, 0.5-0.6 Torr, 60 s) using a PDC-001-HP expanded plasma cleaner (Harrick Plasma, Thermo Fisher Scientific) to improve surface functionality. During cell seeding, we applied 0.5 mL of cell suspension containing approximately 3.94 × 10^5^ cells to each dry scaffold/well, and TCPs were incubated for 15 min to allow good cell attachment. After incubation, we added fresh medium to each well and monitored the cells on the scaffolds every other day for imaging and necrotic core detection.

NIH/3T3-GFP fibroblast growth was monitored using an optical fluorescence microscope (EVOS M5000; Thermo Fisher Scientific) equipped with GFP, DAPI, and RFP filter cubes. To illustrate fibroblast morphology, additional cell visualization was carried out by staining cell nuclei and actin filaments with Hoechst 33258 and rhodamine phalloidin, respectively.

### Cell Viability Assessment

NIH/3T3-GFP fibroblast viability within the 3D scaffolds was evaluated on Day 14 using the resazurin-based PrestoBlue assay. Triplicate samples were studied for each scaffold group. Cells cultured in surface-treated 6-well plates (~3.94 × 10^5^ cells per well) during a 3-day period were used as the positive control, whereas culture medium alone served as the negative control. For the assay, scaffolds were transferred to new 6-well suspension culture plates containing 3 mL of fresh medium per well. PrestoBlue reagent was added at 9 vol % of the total medium volume, and the samples were incubated for 4 h. At the end of the incubation, aliquots (200 μL) were transferred into a 96-well plate, and fluorescence was measured using a multimode microplate reader (Varioskan LUX; Thermo Fisher Scientific) at excitation/emission wavelengths of 560 nm/590 nm.

Statistical analysis was performed to validate the experimental results, with one-way ANOVA used to assess the significance of differences in cell viability among the scaffold groups.

### Cell Counting

Following viability assessment, cell numbers were determined after trypsinization. At least three scaffolds from each group were analyzed. Prior to trypsinization, the outer frames of the touch-spun fiber scaffolds were carefully removed with tweezers, and the samples were transferred to new 6-well suspension culture plates to ensure that only scaffold-associated cells were counted. After medium removal, scaffolds were washed twice with PBS, then 1 mL of trypsin was directly added to each well. Trypsinization was performed for 5 min at 37 °C under 5 % CO_2_. Subsequently, 2 mL PBS was added, and residual cells were detached by vigorous pipetting. The resulting cell suspensions were collected into 15 mL tubes, centrifuged at 1,500 rpm for 5 min, and 100 µL aliquots were used for cell counting with an automated cell counter (Scepter 3.0; Sigma-Aldrich).

## Results and Discussion

In this study, three groups of touch-spun 3D scaffolds were fabricated using a touch-spinning apparatus and a system of grid spacers and frames. Fiber sheets produced during the spinning process were assembled into four-layered 3D scaffolds, with each layer consisting of two orthogonally aligned (crisscrossed) fiber sheets. The scaffold groups differed in interfiber spacing, which was controlled by adjusting the Z-axis speed of the collecting frame within the setup chamber. Optical microscopy analysis revealed that fibers produced by the touch-spinning method exhibited a uniform diameter of 9.08 ± 0.63 μm. The average interfiber spacing for Groups 1, 2, and 3 was 23.24 ± 14.22, 50.61 ± 15.35, and 84.05 ± 14.23 μm (N = 50), respectively. Representative optical microscopy images of microfiber patterns with varying interfiber spacing are shown in **Figure 1a-c**, and the corresponding fiber characteristics are summarized in **Figure 1d**.

**Figure 1.**
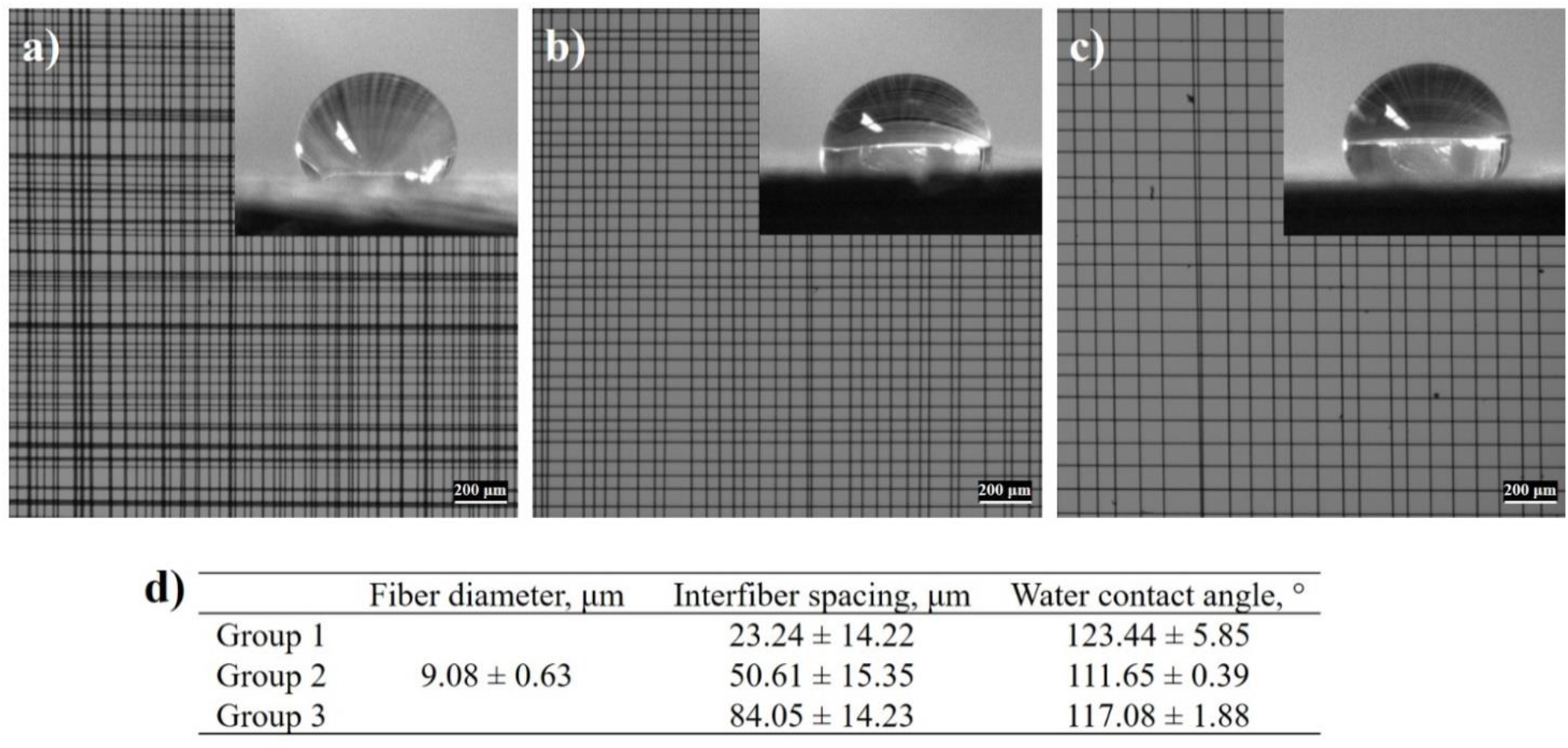
Optical microscopy images (5× magnification) of crisscrossed microfiber sheets with varying interfiber spacing: a) Group 1, b) Group 2, and c) Group 3. Insets show water contact angle measurements used to assess surface wettability; d) characterization of touch-spun microfiber sheets

Although the touch-spinning technique employed in this work enables the production of highly aligned fibers with uniform diameters down to the nanoscale, the uniformity of interfiber spacing was less consistent. The highest precision was achieved for larger spacing values. In addition, slight deviations from the targeted interfiber spacings of 20, 40, and 60 μm were observed.

Water contact angle measurements revealed that all PCL/PEO-based microfiber patterns exhibited hydrophobic surface characteristics, with contact angles exceeding 110°. The relatively low variability within each group indicates consistent fiber surface wettability, while only minor differences were observed among patterns with varying interfiber spacing. NIH/3T3-GFP fibroblasts typically prefer moderately hydrophilic surfaces for optimal adhesion. The water contact angle for a flat PCL coating is 80 ± 5°, while on the fiber mat it approaches 120 ± 5°, reflecting a composite surface effect. This behavior arises because only portions of the water droplet (~ 2 mm in diameter) are in direct contact with the PCL microfibers, whereas most of the droplet interfaces with air.

In this research study, we demonstrate that fibroblasts on flat coatings display sufficient elongation at low confluency, with pseudopodia extended across tens of micrometers, indicating the well-known good affinity of the cells for PCL-based surfaces. Illustrative images of fibroblast morphology on PCL/PEO films (obtained via spin coating onto polystyrene coverslips) after staining for nuclei and actin are presented in **Figure 2**. The incorporation of a fibrous 3D microarchitecture was anticipated to further enhance fibroblast adhesion and proliferation compared with polymer films.

**Figure 2.**
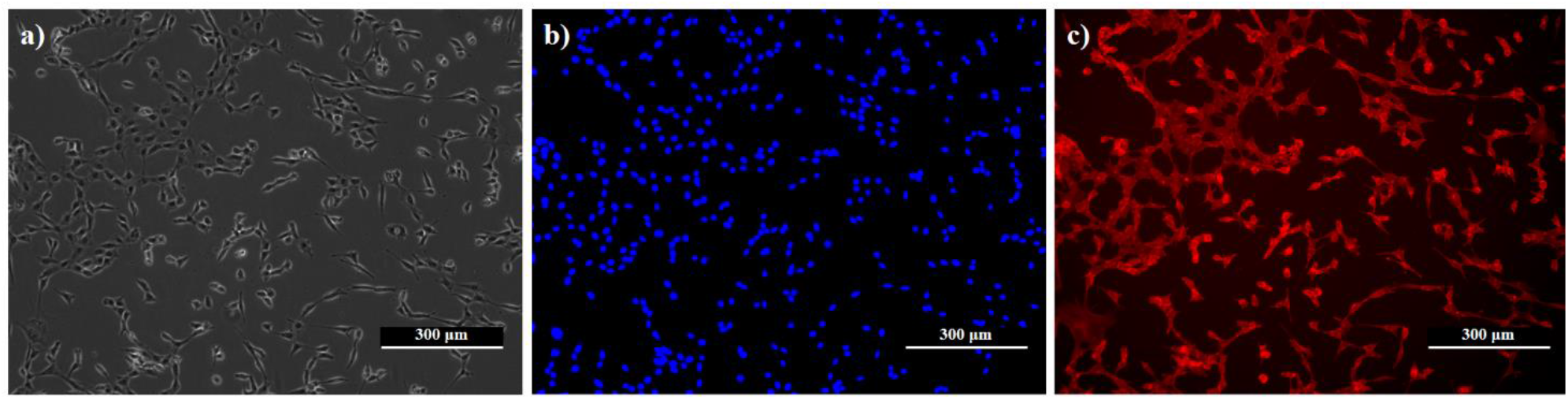
Fluorescence microscopy images (10× magnification) of NIH/3T3-GFP fibroblasts on PCL/PEO films: a) image captured in the TRANS channel showing overall cell morphology, b) nuclei stained and captured in the DAPI channel, and c) actin stained and captured in the RFP channel.

To evaluate scaffold performance across all three groups and determine the optimal interfiber spacing for long-term cell culture, experiments were conducted over a 14-day period. NIH/3T3-GFP fibroblasts were seeded at equal high densities to ensure rapid formation of dense cell cultures within the touch-spun 3D scaffolds. The progression of fibroblast growth within the scaffolds over an 11-day period is shown in **Figure 3**. As cell density increased over time, visualization using standard fluorescence microscopy became challenging, necessitating confocal microscopy for accurate imaging. Thus, by Day 13, distinguishing cell growth between different scaffold layers was no longer feasible in some scaffolds.

**Figure 3.**
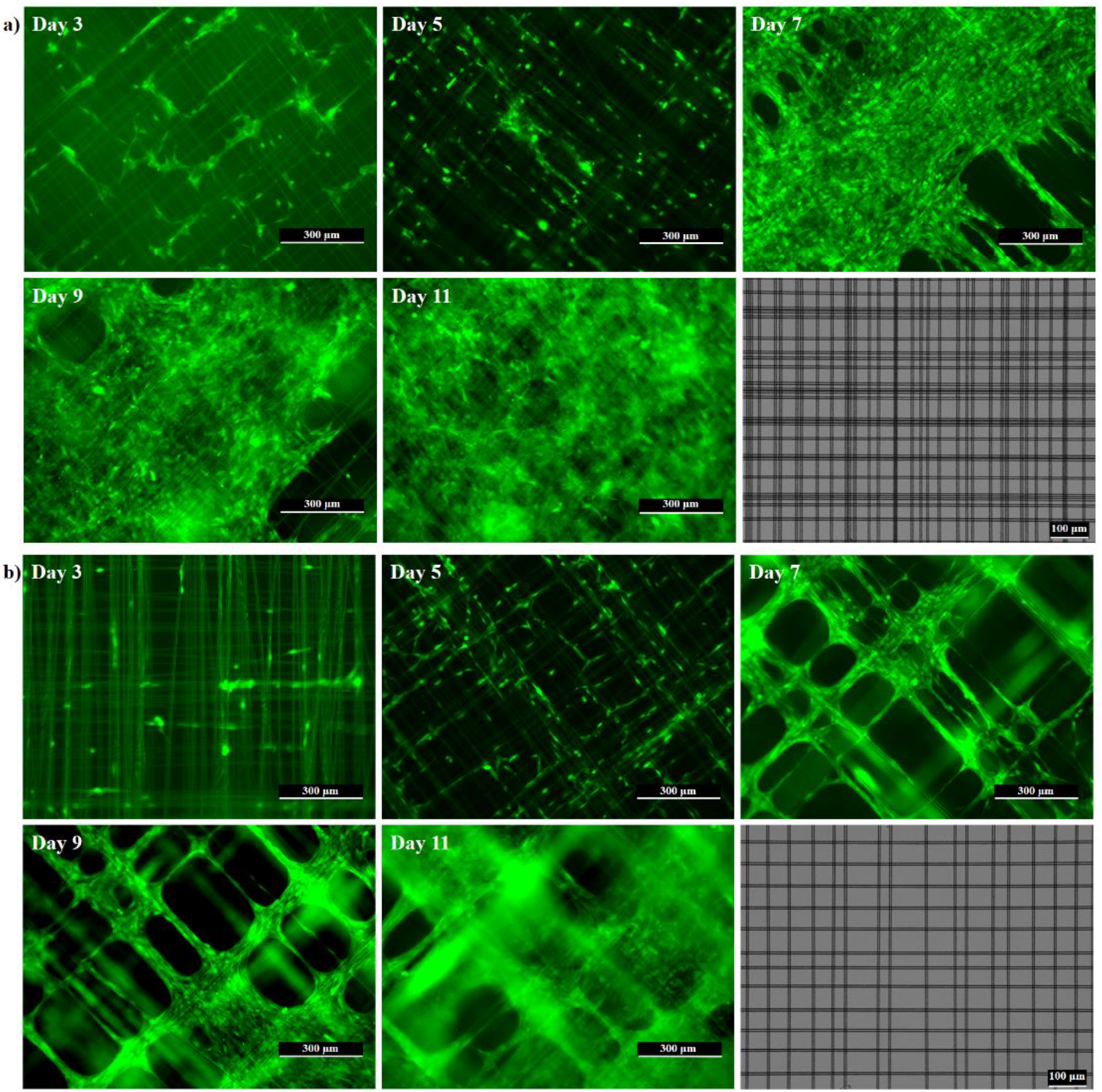
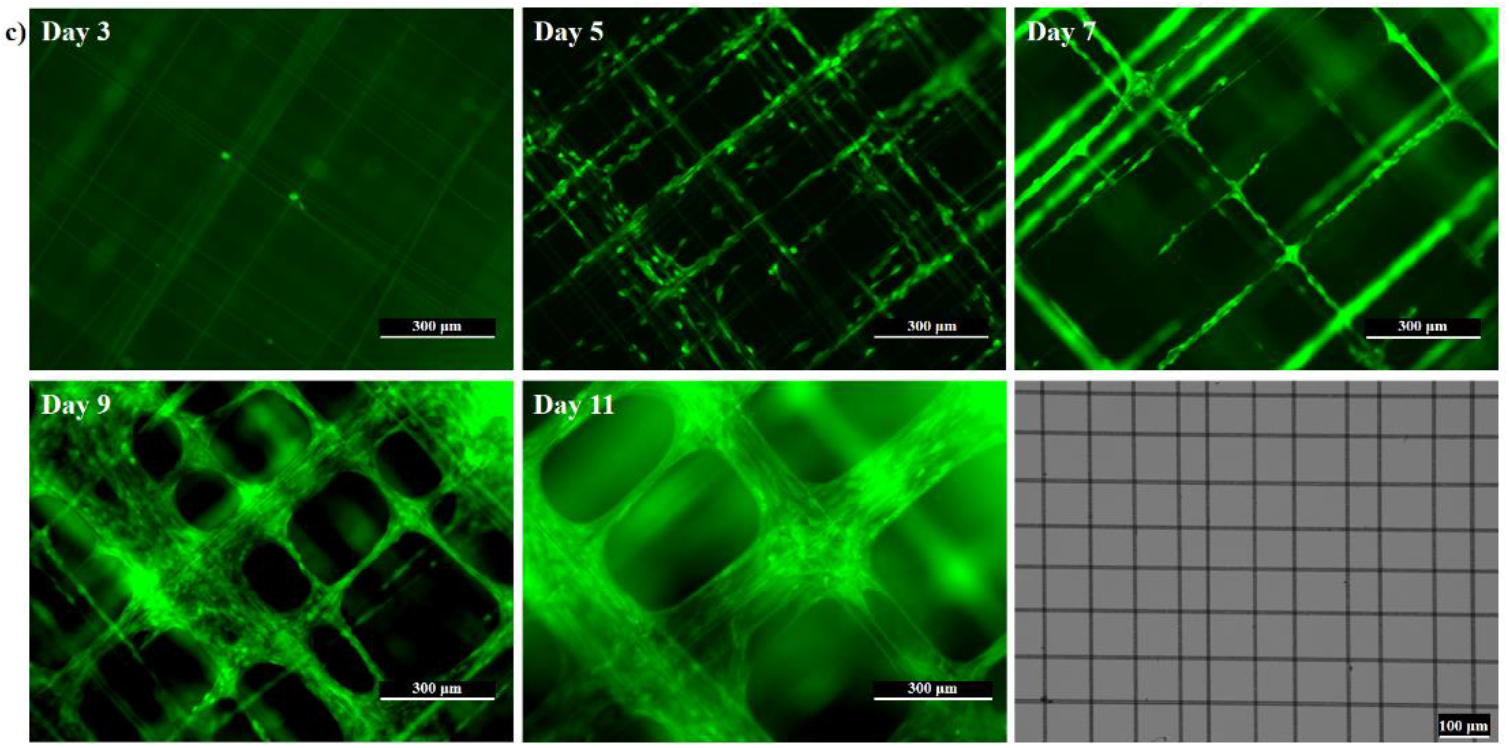
Progression of NIH/3T3-GFP fibroblast growth within touch-spun 3D scaffolds. Fluorescence microscopy images (10× magnification) of cells captured in the GFP channel, alongside optical microscopy images (20× magnification) of individual fiber sheets with varying interfiber spacing: a) Group 1, b) Group 2, and c) Group 3.

Analysis of cell imaging showed that fibroblast growth on aligned microfiber patterns follows a characteristic pathway: cells initially position and elongate along the fibers, and colonization of the square voids within crisscrossed fiber layers begins at the edges, gradually progressing toward the centers. The voids begin to exhibit a circular morphology as they become partially occupied by cells. This process ultimately results in complete void occupancy and the formation of a continuous cell sheet. Additionally, as growth progresses, deviations from the initial fiber alignment become apparent.

By Day 5 of cell culture, no significant differences in fibroblast proliferation were observed among the three groups. However, by Day 7, cell growth within 3D scaffolds with a smaller interfiber spacing of 23.24 ± 14.22 μm intensified markedly, resulting in the occupation of most of the voids and the initiation of dense culture formation. Imaging analysis indicates that fibroblasts reach full confluency within individual layers in scaffolds from Group 1 around Day 11-12 (**Figure 3a**). Nevertheless, layer occupancy is not uniform, with some layers remaining sparsely populated. Once maximal confluency is reached within a layer, cells begin colonizing the interlayer voids. In Group 2, scaffolds with a medium interfiber spacing of 50.61 ± 15.35 μm, show slightly slower cell proliferation, and dense culture formation begins around Day 11 (**Figure 3b**). As further cell culture experiments revealed, fully populated scaffold layers were observed only by Day 16. In scaffolds of Group 3, which exhibited the largest interfiber spacing of 84.05 ± 14.23 μm, complete void occupation requires an even longer period of time (**Figure 3c**).

A slower progression toward full cell confluency within the layers, resulting from larger interfiber spacing, is not necessarily detrimental, as it extends the culture period within the 3D scaffolds, supports healthier cell growth, and reduces the risk of necrotic core formation. Thus, in Group 1, necrotic core formation was observed on Day 14; in contrast, Groups 2 and 3 did not show early signs of cell necrosis. At this time point, cell viability was assessed for all scaffold groups using a non-destructive assay (**Figure 4a**). Further, cell counts were obtained via trypsinization to evaluate scaffold capacity (**Figure 4b**).

**Figure 4.**
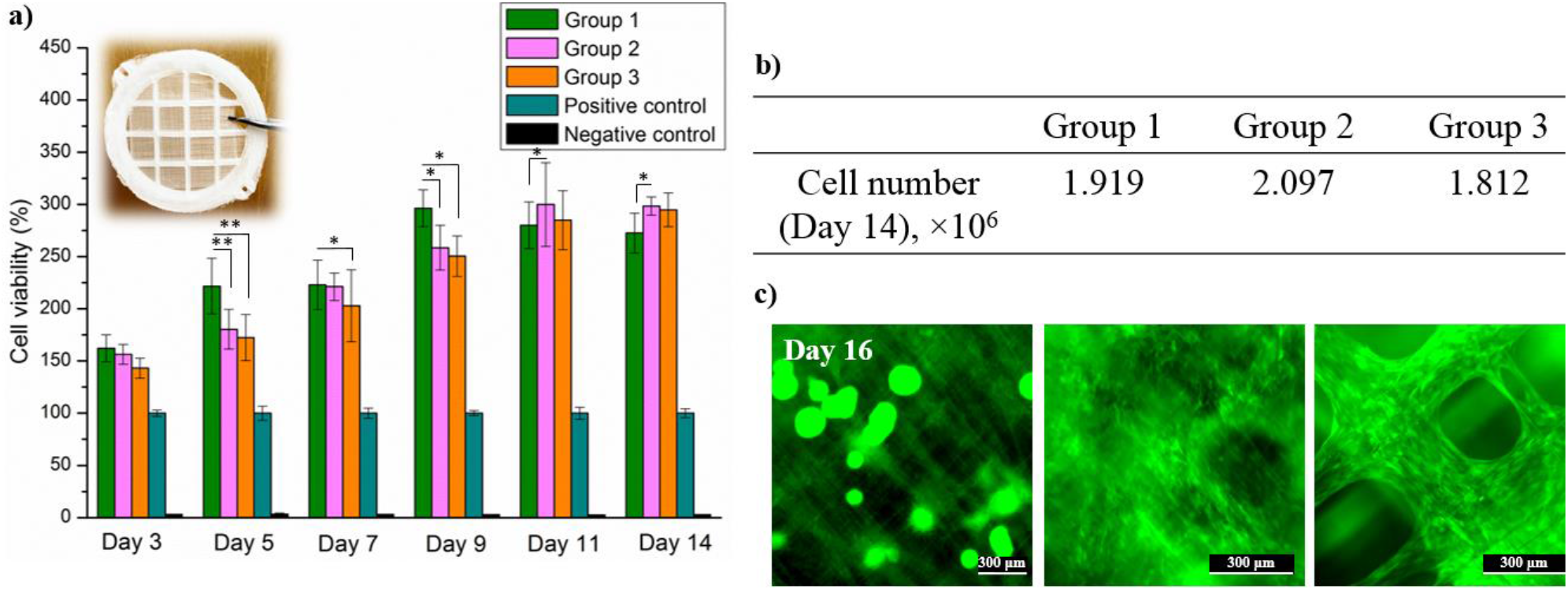
Assessment of scaffold performance: a) PrestoBlue assay outcomes (inset shows a view of the 3D scaffold assembled using the previously reported approach [30]). Statistical analysis was applied to validate the experimental results, with one-way ANOVA used to determine the significance of differences in cell viability; b) cell counting data; and c) representative fluorescence microscopy images of cells on scaffolds captured in the GFP channel on Day 16. From left to right: Group 1, Group 2, and Group 3.

PrestoBlue assay results demonstrated enhanced cell viability in all three scaffold groups compared to the positive control. In this study, the positive control group consisted of cells seeded in empty TCP wells and cultured until maximum confluency was reached. A three-day culture period allowed the cells seeded at the selected density to form a healthy, confluent monolayer in each well, without excessive cell overgrowth that could lead to viability signal reduction. Therefore, this configuration – a confluent cell monolayer formed by Day 3 – was considered to represent the highest viability signal detectable for 2D cell culture under the experimental conditions and was utilized as the positive control. According to the PrestoBlue assay results, no significant difference in cell viability was observed during the early stages of cell culture. Throughout the first five days, Group 1, characterized by the smallest interfiber spacing, exhibited the highest cell viability, significantly exceeding the levels observed for Groups 2 and 3. This trend suggests that a smaller interfiber spacing may facilitate 3T3-GFP fibroblast differentiation by providing closer-spaced biomechanical support points for cell and protein adhesion. The cell viability dominance of Group 1 was maintained through Day 11, at which point a slight reduction in the viability signal occurred. Such a decrease in cell viability may reflect the transition from active cell proliferation toward a confluent state, as the available volume becomes increasingly occupied and the localized regions of overpopulation with fibroblasts appear. At the same time, the viability of cells seeded onto Group 2 and 3 scaffolds consistently increased throughout the culture period, reaching the highest values at the termination of the experiment on Day 14. While no necrotic cores were observed on Day 14, fluorescence microscopy revealed densely populated areas within the touch-spun 3D scaffolds of Group 1, and the cultures were terminated for all groups. The cell culture experiment suggests that the scaffold architectures of Groups 2 and 3 provide a more favorable microenvironment for fibroblast infiltration and proliferation. Despite the lower density of mechanical support points at fiber intersections compared with Group 1, both groups demonstrated a progressive increase in cell viability and ultimately exceeded the levels observed in Group 1. This finding indicates that, while smaller interfiber spacing initially promotes cell attachment and provides enhanced biomechanical cues, a greater spacing between adjacent fibers may facilitate proliferation over longer culture periods.

Cell counting results indicate no significant differences among the three groups, leading to an important conclusion: 3T3-GFP fibroblast growth within the studied 3D scaffolds eventually reaches a saturation level of approximately two million cells across all groups by Day 14, despite the distinct cell viability profiles obtained via PrestoBlue assay. The maximum number of cells harvested from the scaffolds was 2.113 × 10^6^. It is worth noting that in our analysis, living and damaged/dead cells were distinguished using an advanced automated cell counter. Thus, fibroblasts with a diameter of less than or equal to 10 µm were considered to represent cell fragments or non-viable cells and therefore were excluded from the datasets.

As anticipated, a prolonged culture of NIH/3T3-GFP fibroblasts in Group 1 scaffolds led to the formation of necrotic cores shortly after the 14-day period. As illustrated in **Figure 4c** (left image), necrotic cores are aggregates of dead or low-viability cells that form spherical structures, which can reach several hundred micrometers in diameters. Although the majority of cells within these necrotic cores are non-viable, some cells may retain limited viability. Therefore, GFP fluorescence may still be detected during imaging. As the aggregate becomes larger, cells located in the interior experience limited oxygen and nutrient diffusion and insufficient removal of metabolic waste. This leads to reduced metabolic activity, hypoxia, and eventually cell necrosis. The presence of dead cells in the environment affects other living cells (outside the core), as the products of necrosis are being released into the environment. Notably, by Day 16, Groups 2 and 3 continued to exhibit normal cell growth, with no evidence of necrotic core formation. Full confluency within individual scaffold layers was observed in Group 2, whereas scaffolds with the largest interfiber spacing still exhibited unoccupied regions.

## Conclusions

This work provides further evidence of the successful integration of touch-spinning with additive manufacturing for the fabrication of well-organized 3D fibrous scaffolds with uniform fiber diameters and tuned interfiber spacing. Among the three scaffold groups investigated, optimal conditions for enhanced cell proliferation within touch-spun 3D architectures were identified. Specifically, interfiber spacings of 50.61 ± 15.35 and 84.05 ± 14.23 μm were found to support healthy growth of approximately two million NIH/3T3-GFP fibroblasts over a 14-day culture period, with potential for extended cultivation. In contrast, the smaller interfiber spacing of 23.23 ± 14.22 μm promoted rapid cellular overpopulation within scaffold layers, which in turn led to cell necrosis during prolonged culture. The medium and large interfiber spacings examined in this study hold promise for extended culture durations exceeding 14-16 days. Overall, these findings advance the design of touch-spun scaffolds and may guide future studies devoted to the fabrication of ECM-mimicking fibrous scaffolds for clinical applications. Although optimal pore sizes have been discussed for electrospun fiber patterns, corresponding data for well-aligned mechanically drawn fibers remain limited. This research letter aims to address this gap and provide an additional consideration for the engineering of these cell culture platforms.

## Author contributions

K.P., M.A.: Writing original manuscript, Investigation, Data curation. N.S.Y., S.M. ^a,b^, M.P.: Methodology, Investigation, Data curation. S.M. ^a,b,c,*^, V.R.: Supervision, Project administration, Editing.

## Acknowledgements

This research was supported by the Georgia Research Alliance (Grant Title: BioScaffold: Nanofiber 3D Scaffolding Devices for 3D Cell Culture), the USDA National Institute of Food and Agriculture STTR Program, USA (Grant No.: 2023-51402-39329), and the Good Food Institute (Grant Title: Tailored, Edible, and Scalable 3D Fiber Scaffolds for Shrimp Cell Expansion). M.A. was supported by the U.S. Department of Energy, Office of Science, Office of Biological and Environmental Research program under Award No. DE-SC0023338.

## Data availability

All data obtained in this study are fully reported in this published research letter.

